# Quantitative analysis of *C. elegans* transcripts by Nanopore direct-cDNA sequencing reveals terminal hairpins in non trans-spliced mRNAs

**DOI:** 10.1101/2022.04.14.488332

**Authors:** Florian Bernard, Delphine Dargère, Oded Rechavi, Denis Dupuy

## Abstract

Nematode mRNA processing involves a trans-splicing step through which a 21nt sequence from a snRNP replaces the original 5’ end of the primary transcript. It has long been held that 70% of *C. elegans* mRNAs are submitted to trans-splicing. Our recent work suggested that the mechanism is more pervasive but not fully captured by mainstream transcriptome sequencing methods.

In this study, we used Oxford Nanopore’s long-read amplification-free sequencing technology to perform a comprehensive analysis of trans-splicing in worms.

We demonstrated that spliced leader (SL) sequences presence at the 5’ end of the messengers affected library preparation and generated sequencing artefacts due to their self-complementarity. Consistent with our previous observations, we found evidence of trans-splicing for most genes. However, a subset of genes appears to be only marginally trans-spliced. These messengers all share the capacity to generate a 5’ terminal hairpin structure mimicking the SL structure providing a mechanistic explanation for their non conformity. Altogether, our data provides the most comprehensive quantitative analysis of SL usage to date in *C. elegans*.

## Introduction

Trans-splicing is an RNA maturation process originally discovered in Trypanosomes, where gene expression is polycistronic and the resulting long RNAs needs to be matured into monocistronic units before being fully functional. In these parasitic organisms the first Spliced Leader sequence (SL) was detected during the characterization of cDNAs clones encoding surface glycoproteins (VSGs)^1^. It has then been shown that coupling of trans-splicing and polyadenylation is a crucial step of the maturation of the RNAs in Trypanosomes^2^. In this protists genes do not contains introns, therefore the spliceosome sole function is to perform trans-splicing so that all the mRNAs begin with a SL sequence of 39nt derived from the 5’ end of a 137nt RNA called medRNA. After the discovery of trans-splicing in trypanosomes, it was found to exist in other metazoans, including cnidarians, ctenophores, rotifers, flatworms, nematodes, crustaceans and sponges. However, trans-splicing has not been found in any plants, fungi, insects or vertebrates^3^.

In the nematode *Caenorhabditis elegans*, trans-splicing was initially uncovered during the study of the 5’ extremity of the actin mRNA, which was found to contain an additional 22nt sequence that was absent from the genomic copy of the gene^4^. This first nematode SL sequence (SL1) was revealed to be donated by a 100nt small nuclear ribonucleoprotein particle (snRNP). This process is closely related to cis-splicing (intron removal) where the 5’ splice site is on the SL RNA, and the site of SL addition (3’ splice site) is on the pre-mRNA^5^. The reaction happens through a branched intermediate, similar to the lariat of cis-splicing: cleavage of the SL sequence by the 5’ region of the pre-mRNA in a first step, leading to the formation of a branched intermediate between the two, and splicing of the SL to the first exon of the pre-mRNA in a second step. The excised 5’ region of the pre-mRNA is called the outron^6^.

In 1989, a second SL sequence was found. This SL2 sequence is the same size as SL1, however with a different sequence^7^. Since the original discovery, many more mRNAs have been found associated with SL2 sequences and many operon-like clusters have been identified^8,9^. A whole-genome microarray analysis has demonstrated a robust correlation between gene clusters and SL2-containing genes^10, 11^. In this study, the authors have identified more than 1,000 clusters for which downstream messengers are trans-spliced to SL2, yet it remained a mystery how genomic position can affect trans-splicing specificity. It was hypothesized that gene clusters acts in a similar way as bacterial operons, with the transcription of the entire cluster by a promoter situated on the 5’ extremity of the region. However, while in bacteria mRNAs from operons remain polycistronic, in *C. elegans* they are processed into monocistronic units in a similar fashion as to how trans-splicing and polyadenylation are coupled in trypanosomes^12^.

Early global analyses of trans-splicing estimated that 70% of *C. elegans* messengers are trans spliced^9, 13^. A meta-analysis of RNA-seq datasets amended this number to 85% and confirmed a trend that the least expressed genes had the most propensity for not being trans-spliced^14^. This was suggestive of a lack of detection of trans-splicing for rare mRNAs rather than a distinct subset of genes bypassing the generic mRNA maturation process. This prompted us to hypothesise that the actual fraction of trans-spliced genes could be higher still. To test this hypothesis we decided to apply a different RNA-seq method to see wether it could improve our characterization of 5’ extremities.

In 2015 by Oxford Nanopore Technologies commercialised a new sequencing method relying on the ability of a molecule to affect ionic currents based on the amount of space it takes inside a nanopore, therefore making it is possible to reconstruct the sequence of nucleic acids being translocated through a membrane by measuring changes affecting the current^15,16^. In 2020, two teams have performed transcriptome-wide analysis using Nanopore direct-RNA sequencing in *C. elegans*. The authors have demonstrated that full-length reads could be used for the easy detection of novel splice isoforms. However this technology does not provide a good enough read out for studying trans-splicing events^17,18^, as shown by a previous study comparing direct-RNA and direct-cDNA sequencing which reported that direct-RNA reads exhibits shorter 5’ extremities compared to reads generated with direct-cDNA sequencing^19^. We therefore selected direct-cDNA sequencing as our experimental strategy for amplification-free quantitative analysis of spliced leader sequences on the 5’ extremity of *C. elegans* mRNAs. Our experiments first revealed how the presence of Spliced Leader sequences affects the Nanopore library preparation process. Our data provides the most comprehensive quantitative analysis of SL usage to date in *C. elegans* and allowed us to formulate an hypothesis regarding a common feature of non trans-spliced messengers.

## Results

### *C. elegans* Spliced Leaders interfere with direct-cDNA library preparation

To comprehensively describe the trans-splicing repertoire of *C. elegans* nematodes we selected direct cDNA sequencing. After performing three independent experiments and mapping our reads onto *C. elegans* genome, we controlled their correct alignment by looking at them in Integrative Genome Viewer (IGV), and noticed an unexpected, reproducible strand bias in favor of antisense reads (Fig. 1a). Out of ∼1,3 million reads obtained in these initial experiments, 95% were antisense reads. Additionally, we observed that these antisense reads harbored extended 5’ soft clipped region. The soft clipped region is the part of the sequence that is beyond the portion of the read that is aligned to the genome (Sup. Fig. 1). In our case the soft clips are expected to contain the sequences of oligonucoleotide linkers used in the sequencing library preparation as well as spliced leader sequences on the 5’ end, and polyA tail on the 3’ end. Indeed, that was the case for the minority of sense strand reads (Sup. Fig. 2). On the other hand, the 5’ soft clipped part of the antisense reads had a median size of 416bp, well beyond what is expected at the 5’ end of the cDNA produced by the Nanopore library preparation protocol (Sup. Fig. 3). We investigated the content of these extended unaligned sequences and noticed that ∼33% of these contained a supplementary alignment matching to the sense strand of the original alignment (Sup. Fig. 4). These supplementary alignments were of lower quality than the primary ones and we wondered if this could be the result of poor sequence quality in the long soft-clipped region. When plotting the base-quality of a single read, we observed the quality was highly variable along the sequence. To mitigate this effect, we plotted the average base-quality of groups of one million reads and then computed the mean Q-score at every position centred around the 5’ end of the alignment. This confirmed that the base quality of the long soft clips is significantly lower than that of the aligned portion of the reads (Fig. 1b & Sup. Fig. 5).

**Fig. 1:**
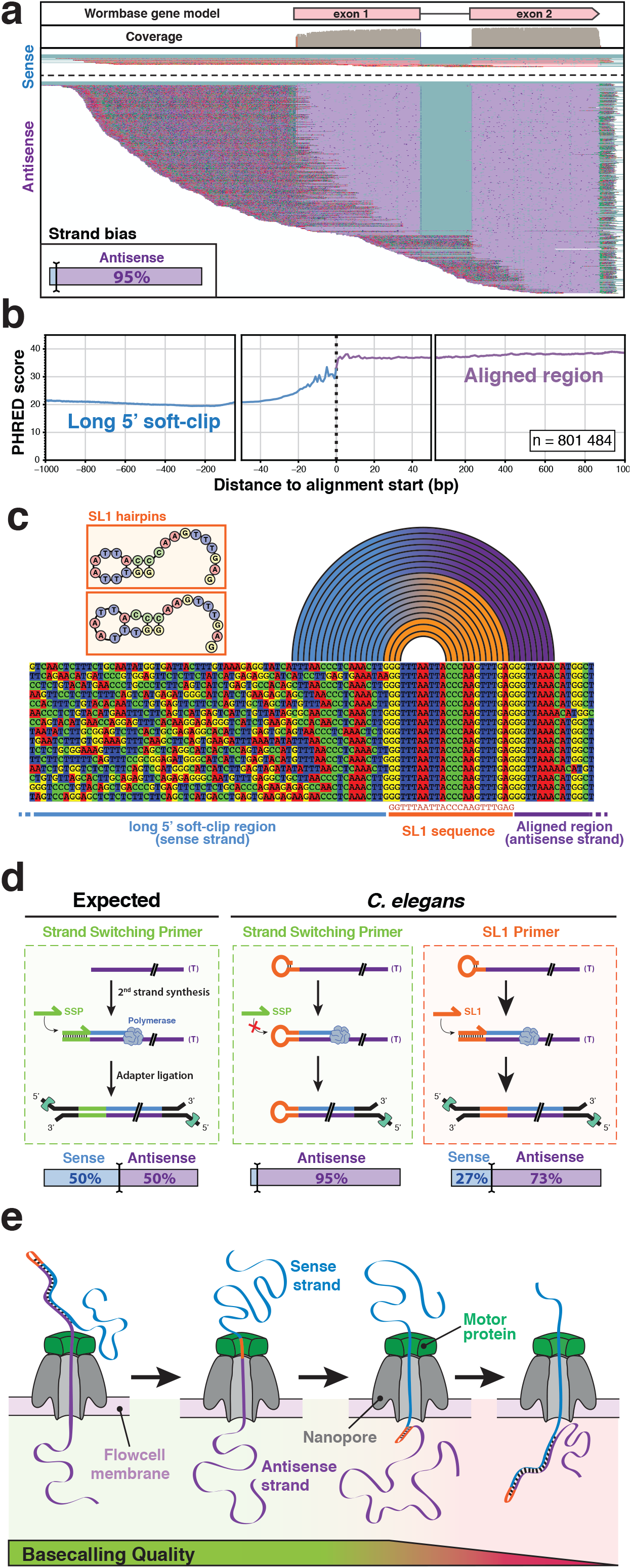
Strand bias in Nanopore direct c D N A sequencing of *C. elegans* transcripts. **a** Typical genome browser (IGV) view of direct cDNA reads aligned on a *C. elegans* gene (here gene *lys-1*). Aligned bases from the sense strand reads are shown in pink and aligned bases from the antisense strands in purple. Unaligned soft clip region is shown with mismatches colored according to the observed base. Inset: overall strand bias observed across all detected transcripts. **b** Base quality measured in 5’ soft-clip and primary alignments. The average base quality value over ∼800,000 individual Nanopore reads was measured. **c** Typical alignment of reads at the interface between the transcript match and the long soft-clipped region. The first bases of the soft-clip commonly correspond to the SL1 sequence followed by a partial antisense SL1 segment (blue to orange arcs) consistent with the extension of one of the endogenous hairpins represented as insets. The arcs indicate base pairing between the 5’ soft clip and the trans spliced antisense strand. **d** Schematic model of molecular events occurring during the second strand synthesis step in various conditions of direct cDNA sequencing library preparation. Bottom : observed strand bias measured after Minion sequencing. **e** Schematic representation of the progression of a hairpin cDNA substrate through the pore during Nanopore sequencing. The helicase activity of the motor protein maintains a steady rate of transfer for the antisense strand while unwinding the double stranded cDNA. For the second strand the absence of double helix on the input side and/or the kinetics of the base pairing on the other side of the pore affects the signal quality and prevents accurate base calling of the sense strand.

The observation of an overwhelming bias for antisense reads combined with the presence of a low quality extended 5’ soft-clip that could be partially mapped to the cognate sense strand, was indicative of the systematic generation of long hairpin cDNAs. Since this behavior was not reported by other users of the Nanopore direct cDNA sequencing kit we hypothesised that the origin of this phenomenon may be linked to the widespread addition of spliced leaders at the 5’ ends of the transcripts through trans splicing. To test this, we selected and aligned full-length reads that displayed a complete SL1 sequence at the beginning of their soft clipped sequence (Fig. 1c). Looking at these reads where the sequence quality next to the alignment region was sufficient to identify the neighboring nucleotides we could confirm that in these sequences the two guanine bases at the 5’ extrimity of the SL1 sequence had paired with two of the three cytosines in the middle of the spliced leader leading to self-priming of long hairpin cDNAs during the second strand synthesis step of the library construction. The left panel of Figure 1d illustrates the normal second strand synthesis step as described in Oxford Nanopore documentation: a strand switching primer hybridizes to the terminal cytosines added by the terminal nucleotidyl transferase activity of the reverse transcriptase on the first strand (antisense) cDNA and initiates the synthesis of the sense strand providing the double stranded substrate for the adapter ligation that will lead to the generation of an equal proportion of sense and antisense reads during the Nanopore sequencing^20, 21^. In our *C. elegans* samples the presence of a self-complementary splice leader at the 5’ end of the mRNAs mostly prevents the entry of the strand switching primer (SSP) and leads to the strand bias we observed. To test this model, we performed additional library preparations replacing the supplied “strand switching primer” by a SL1 oligonucleotide. In this configuration, we could partially prevent the formation of the hairpin and recover 40% of sense strand reads (Fig. 1d, right panel) as well as the corresponding fraction of antisense reads with a short soft clipped region (not shown). We also attempted a direct cDNA library preparation in the absence of SSP and confirmed it was not required for effective Nanopore sequencing of *C. elegans* transcripts (see Table 1 and Sup. Fig. 6). The obtained strand bias was comparable to that observed in presence of SSP. Thus, we concluded that the presence of a spliced leader sequence (either SL1 or SL2) at the 5’ end of *C. elegans* mRNAs generates double stranded hairpins of which only the first (antisense) strand can be effectively read by Oxford Nanopore sequencing. Indeed, the helicase currently used is calibrated to provide constant speed of passage of each individual molecule through the channel by unwinding a double-stranded cDNA molecule. In our case, when the first strand is fully processed the second strand is now single-stranded on the helicase side and is (likely) re-annealing on the other side of the membrane. In this configuration, the speed of passage through the pore becomes too perturbed to provide good quality sequence information for the second strand strand and thus preventing accurate base calling for a fraction of SL sequences and most of the length of the second strand (Fig. 1e, from personal communications with ONT support team).

**Table 1:** Minion Direct cDNA runs included in this study. SSP: Strand Switching Primer, SL1: SL1 primer, NP: No primer

| Exp name | Primer for 2nd strand synthesis | Transcriptome mapping |  |  | Antisense strand |
| --- | --- | --- | --- | --- | --- |
|  |  | Alignments | Genes | Isoforms |  |
| SSP_1 | SSP | 856 699 | 12 661 | 19 079 | 98.2% |
| SSP_2 |  | 265 555 | 12 586 | 18 351 | 97.6% |
| SSP_3 |  | 215 150 | 11 790 | 17 426 | 93.0% |
| SSP_4 |  | 329 046 | 15 078 | 22 808 | 93.8% |
| SSP_5 |  | 75 874 | 9 858 | 13 416 | 96.3% |
| SSP_6 |  | 520 347 | 12 558 | 19 710 | 92.3% |
| SL1_1 | SL1 | 6 015 856 | 15 080 | 24 474 | 73.3% |
| NP_1 | None | 106 081 | 6 736 | 8 914 | 98.2% |
| NP_2 |  | 1 962 645 | 13 041 | 20 486 | 98.3% |
| NP_3 |  | 398 165 | 9 781 | 14 107 | 98.6% |
| NP_4 |  | 54 418 | 5 343 | 6 890 | 98.3% |
| NP_5 |  | 353 154 | 9 681 | 14 298 | 98.4% |

### A significant fraction of SL2 trans-spliced genes are not part of an operon

*C. elegans* spliced leader precursors are expressed from 30 individual trans <u>S</u>pliced <u>L</u>eader <u>S</u>equence (*sls)* genes. SL1 spliced leaders are expressed from 12 genes (*sls-1* to *12*) interspersed in the rRNA cluster located on chromosome V. There are 11 distinct SL2 variants that are produced from 18 *sls-2* genes located on chromosomes I to IV (Fig. 2a). From ∼11 millions reads collected in our direct cDNA sequencing, we extracted ∼2.8 million reads where the soft clipped sequence could be unambiguously attributed to a specific spliced leader to determine the usage frequency of each. The overwhelming majority (86%) of these reads contain the SL1 sequence, as previously reported^13, 14, 22^. Direct cDNA sequencing allowed us to measure the relative usage of each SL2 variant. SL2.1, produced by four genes, is the most prominent, representing ∼20% of all SL2 reads followed by SL2.8 (17%) coming from a single gene (Fig. 2b).

**Fig. 2:**
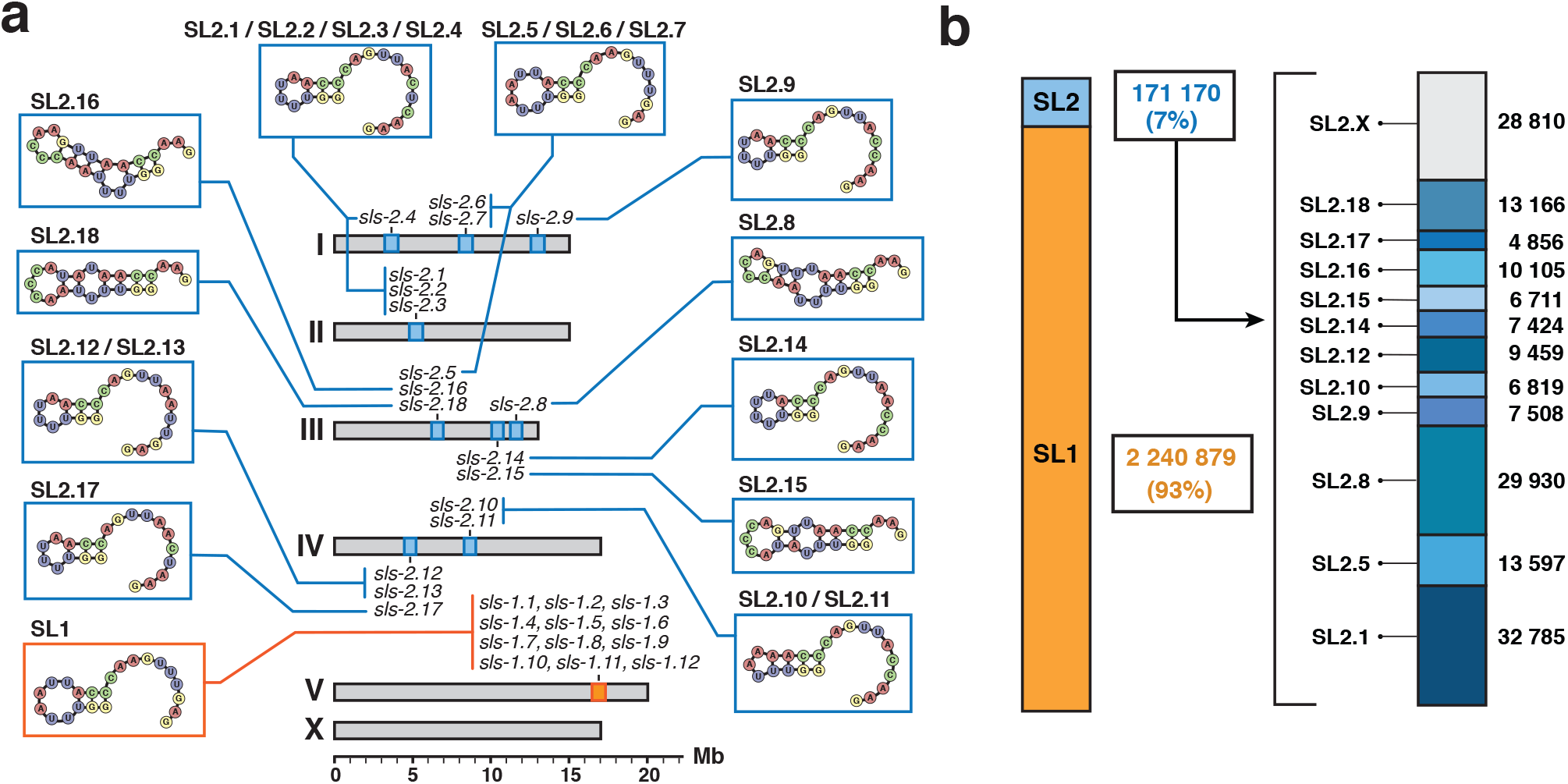
Spliced leaders usage frequency. **a** Genomic location of all *C. elegans sls* genes and structure of their 5’ hairpin strand. **b** Quantification of reads with high quality splice leader sequence reveals the usage frequency of each of the *sls* genes. SL2.x indicates reads for which the specific SL2 variant could not be unambiguously identified.

For each of the trans splicing sites we counted the number reads that could be unambiguously discriminated between SL1 and SL2. For positions with at least 10 such reads, we plotted the ratio of SL2/SL1 in relation to the distance to the closest annotated upstream coding gene (Fig. 3a). We confirmed previous observations that trans splicing acceptor sites are strongly favoured by one SL sequence or the other and that SL2 trans-splicing is strongly preferred when the nearest gene is located ∼100 nucleotides upstream of the trans splicing acceptor site^13, 14^. However, our data demonstrate that this operon-like genome organisation does not fully explain all instances of SL2 proclivity. While 700 out of 1011 genes are located within 200 pb downstream of the closest upstream coding gene a significant number of SL2 genes are several kb downstream of their closest neighbor. We investigated the possible presence of cryptic termination signals^23^ in the upstream region of isolated SL2 genes but didn’t find evidence supporting that explanation (not shown).

**Fig. 3:**
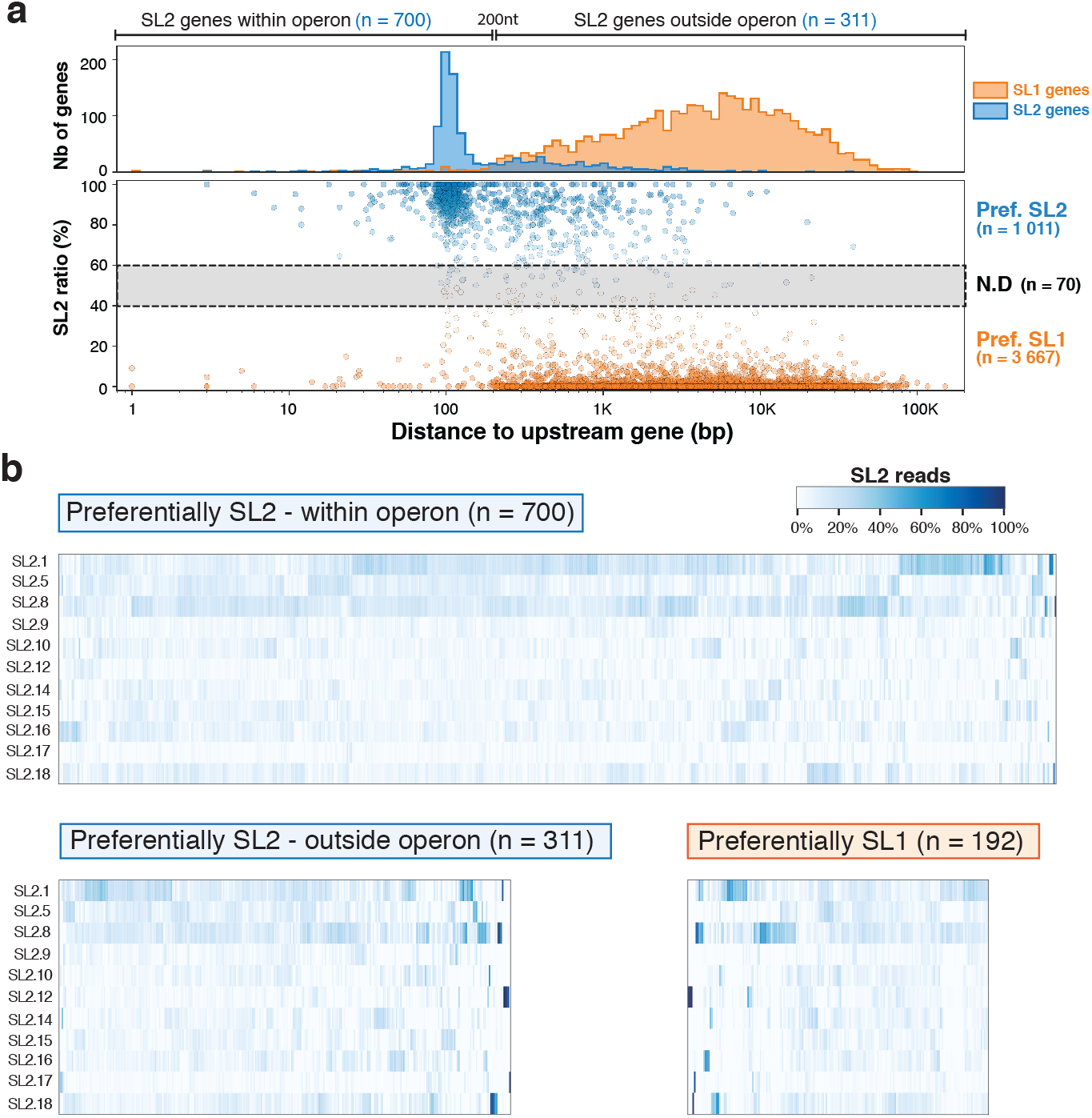
Quantitative analysis of SL sequence selectivity. **a** bottom: A scatter-plot of the ratio of SL2/SL1 reads found against the distance to the closest upstream gene. Positions with at least a 60% bias are colored red for SL2 and blue for SL1. Top: Distribution SL1 and SL2 biased positions confirms previous observations of a strong preference for SL2 splicing to occur at a distance of ∼100nt downstream from the preceding gene. **b** In each column of the heatmap we represent the frequency of usage of each SL2 variant for each SL2 trans spliced site represented in a. Top: Genes located less than 200nt downstream of another coding gene. Bottom: Genes located further away from the closest upstream coding gene with genes predominantly SL2 on the left and predominantly SL1 on the right.

We next measured the proportion of SL2 variants found for each of the majoritarily SL2 genes. The distribution showed no evidence of enrichment for any particular SL2 variant supporting a functional redundancy between all of them (Fig. 3b, see read counts in Sup. Table 2).

### Non trans-spliced transcripts display a terminal hairpin that mimics the structure of the spliced leaders

Our previous meta-analysis of a compendium of RNA-seq data generated by Illumina sequencing indicated a strong positive correlation between the level of expression of a given gene and our capacity to detect a trans splicing event^14^. Here we confirmed this correlation between the expression level observed by direct cDNA sequencing and our capacity to find evidence of trans splicing either in our previous meta analysis or this work or both (Fig. 4a). This correlation seems to indicate that trans-splicing is ubiquitous for *C. elegans* transcripts, which contradicts early observations of non trans-spliced messengers. We therefore decided to look more quantitatively at the number of reads with direct evidence of trans-splicing (*i*.*e*. reads where the sequence spans the splice site and at least part of the spliced leader) relative to the total number of full-length reads for a given gene. For each read alignment, we extracted both start and end positions, corresponding respectively to the 5’ and 3’ ending position of the cDNA sequence mapped onto the reference genome. For most genes, we observed a dominant start position coinciding with the annotated gene start, which was indicative of full-length reads. For most genes there is a strong positive correlation between the number of full-length reads and the number of reads with detectable SL, which indicates that most genes overwhelmingly produces full length cDNA reads containing a spliced leader sequence (Fig. 4b). There is however a subset of genes that display significantly less evidence of trans-splicing than the majority, and these include genes that were originally described as “non trans-spliced”. Those genes however still showed the same strand bias as trans spliced genes. We therefore investigated these sequences to identify the origin of the underlying hairpins. In all cases, we found that the 5’ end of the messenger had the capacity to generate a hairpin by local self complementarity (Fig. 4c). For example, the major starting position for full-length transcript for *vit-2* produces two distinct hairpins that we could identify with 7 and 6 base pairing respectively.

**Fig. 4:**
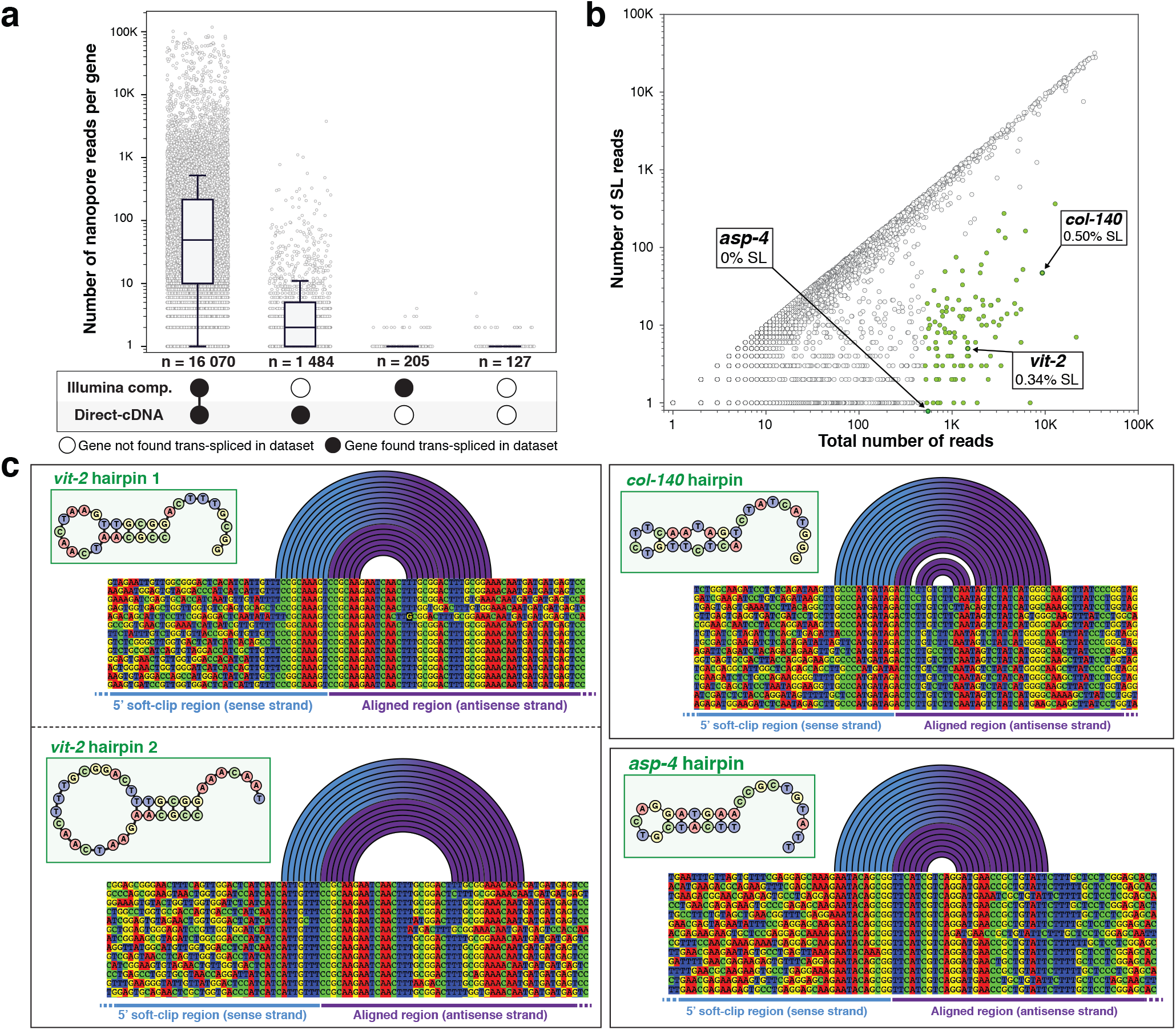
Nature of non- trans-spliced messengers. **a** Gene expression and trans-splicing status. *C. elegans* genes where categorized according to the detection of evidence of trans-splicing in this study and/ or in the compendium of RNA-seq we previously analysed14. For each gene we plotted the number of Nanopore reads as a proxy of expression level. **b** Trans-splicing detection level. For each gene we identified the most frequently detected 5’ alignment starting position and plotted the number of reads at this position vs the number of those reads that for which a Spliced Leader sequence could be detected. Genes for which the most represented start position had at least 100 reads but had less than 10% SL detection are represented in green. **c** Poorly trans-spliced mRNA have a propensity to form a 5’ stem loop structure. For the three genes highlighted in panel b we show a partial alignment of the end of the aligned region and the beginning of the soft clip. Arcs represent base paring.

### Quantitative visualization of Spliced Leaders usage

In the course of this analysis we observed that for a large number of reads the sequence of the hairpin structure was not accessible. We suspected it was due to variability in the individual sequence quality: indeed, when the read quality is lower the alignment may not reach the actual extremity of the antisense cDNA and thus we may not have access to the sequence of the hairpin

itself. We plotted the average sequence quality along three sets of 250,000 reads for which we could identify either a Spliced leader sequence, an endogenous hairpin or neither, and found the latter tend to have lower overall sequence quality when compared to the two other categories (Fig 5a). This lower average quality compared to the two others was also visible in the aligned region of the reads which could explain why, at the proximity of the hairpin structure, the sequence quality did not allow to resolve the end of the first strand. The high quality portion of SL reads extends further than that of endogenous hairpin reads because the splice leader extends the double stranded region by almost 20 nt (see Fig. 1c).

**Fig. 5:**
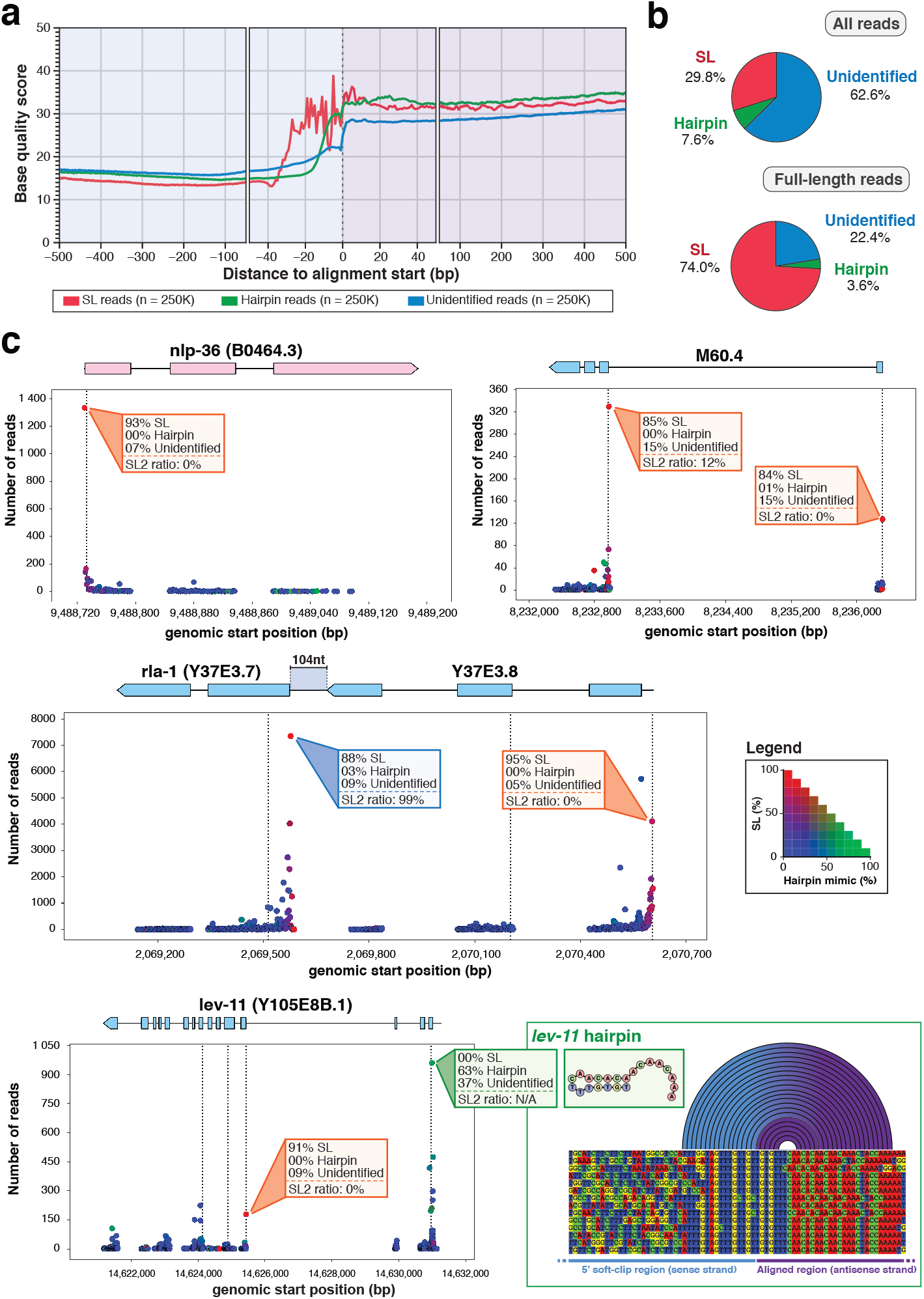
Quantitative representation of trans-splicing prevalence. **a** We randomly selected 250,000 reads from each category and plotted the average quality score around the end of the alignment start position. **b** Repartition of reads for which a trans splicing event could be confirmed (SL), with terminal self complementarity (hairpin) or with unknown 5’ status. **c** Schematic representation of the trans-splicing information recovered in this work. Each alignment start position observed was plotted at the corresponding genomic position with the number of supporting reads. The dots are colored according to the legend on the right, with red indicating a majority of SL reads, green a majority of endogenous hairpin reads and blue reads with no evidence for either. Dotted vertical lines indicate the position of a putative start codon. *nlp-36* displayed a single SL1 trans-spliced site. M60.4 presents two trans-splicing sites indicative of two alternative promoters with distinct SL1/2 ratios. The Y37E3.7/Y37E3.8 operon with typical SL1 and SL2 preference. *lev-11* gene distal promoter carries an endogenous hairpin while the proximal promoter is mainly SL1.

**Fig. 6:**
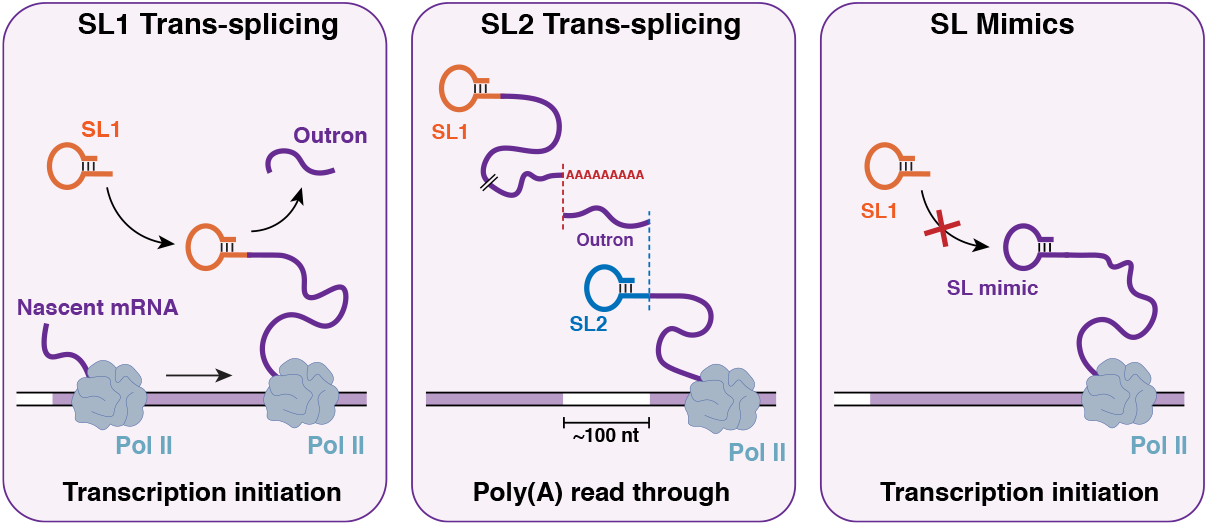
Three types of *C. elegans* mRNA 5’ extremities. mRNAs derived from genes adjacent to their promoters receive mostly SL1. SL2 is preferentially trans-spliced to genes located ∼100nt downstream of another gene. Non-trans-spliced mRNAs have 5’ self-complementarity that mimics the structure of SLs.

Additionally, we observed that the fraction of reads with unidentified hairpin sequence was lower for full-length reads (22.4%) than for all reads (62.6%). Conversely, full-length reads also display a significantly higher proportion of detectable SL sequences : from 29.8% among all reads to 74% among the “full length” reads (see Fig. 5b). Together these observations suggest that reads of lower quality do not allow a complete mapping of their 5’ sequence which precludes the identification of the hairpin that causes the quasi systematic strand bias and the presence of a long “soft-clipped” unaligned region.

We systematically analyzed all start positions for every transcript identified in our direct cDNA sequencing runs. For each gene we generated a visual representation of the identified messenger starts with quantitative information on the usage of each start site and its trans-splicing status see section on Data availability). Fig. 5c presents several examples of such representation. For the gene *lev-11* which features two promoters with different trans-splicing modes. The distal promoter is the most active but seems to produce almost exclusively messengers with endogenous hairpins (99% of reads of sufficient quality to determine the 5’ end structure showed an endogenous hairpin) while the proximal promoter that drives the expression of shorter isoforms displays a systematic use of trans splicing.

## Discussion

In this work, we aimed to provide quantitative information on the 5’ ends of *C. elegans* messengers through the analysis of full-length sequences. The application of Oxford Nanopore direct-cDNA sequencing strategy to *C. elegans* transcriptome revealed that the presence of Spliced Leaders with 5’ self complementarity causes the self priming of the cDNA second strand synthesis during library preparation. While this artefact can be partially circumvented by replacing the regular Strand Switching Primer by a SL1 primer, we did not deem this necessary as the obtained sequences are generally informative up to the end of the antisense strand and a few nucleotides beyond.

Apart from the technical considerations, our observations raise the question of the functional significance of the presence of this stem-loop structure at the extremity of all *C. elegans* SL variants. It has been previously observed that potential stem-loop structures adjacent to the cap are widespread in the spliced leaders of phylogenetically divergent species although the sequence of the self complementarity itself is not conserved^24^. The strong impact we detected on reverse transcribed antisense DNA indicates a strong propensity for this hairpin formation *in vitro* which could be a reflection of a similar structure being typical of worm mRNAs. In the cellular context, such a feature could be an important determinant for the binding of proteins involved either in nuclear export or translation initiation. This hypothesis in reinforced by our discovery that non-trans-spliced genes in *C. elegans* also harbor a 5’ terminal hairpin structure. Additionally, while all Spliced Leaders present a modified Guanine in 5’, endogenous terminal hairpins do not, which would make them uncapped mRNAs^24^. It has been suggested that translational efficiency of “non trans-spliced *C. elegans* mRNA” is lower than for messengers with SLs^25^. Our data indicate that these genes are in fact producing a small fraction of trans-spliced mRNAs while the majority of theis transcripts bears a terminal hairpin that mimics the structure of SLs. It remains to be tested if this reported lower translation rate is due to the suboptimal recruitment of those endogenous hairpins to the translation initiation complex by other protein factors or if the their translation is entirely dependent on the small fraction of those mRNAs that have indeed received a Spliced Leader. More detailed studies will be needed as each endogenous hairpin may differently affect both the capacity of its pre-mRNA to be trans-spliced and its eventual access to the ribosomes if it is not.

Our analysis revealed additional complexity in the *C. elegans* dual trans-splicing system. The majority of SL2 trans-splicing is predominantly targeted to genes located in close downstream proximity to another one, supporting a mechanistic link between mRNA 3’ end formation and the downstream addition of SL2 for resolving multicistronic primary transcripts^26, 27^. Our data however indicate that there is a significant number of SL2 genes that are derived from isolated genes with their own promoter.

## Methods

### *C. elegans* strains and maintenance

Standard protocols were used for maintenance of *C. elegans*. Bristol N2 strain was used as wild-type. Nematodes were maintained at 21°c on Nematode Growth Media (NGM) plates seeded with *E. coli* OP50 as a source of food^28^.

### Synchronization of worms

Worms were grown on 10cm NGM plates in order to obtain adults gravid worms. We recovered the worms by washing the plates with M9 buffer and then proceeded to perform a bleaching following the established protocol^29^. Upon recovery of the eggs, we let them hatch overnight in M9 buffer deprived of a source of food in order to trigger a developmental arrest at the L1 stage. The larvae were then put back to growth on 30cm seeded NGM plates until most of the population was comprised of young adult worms.

### RNA extraction and poly(A) isolations

In order to remove any bacterial contaminants, synchronized young adults were recovered from their plates with clean H20 and washed several times. Following centrifugation (400g, 1min), the supernatant was replaced with clean H20 and the washing step was repeated (at least three times in total). Upon obtention of a clean worm pellet, worms were resuspended in RNA Blue (Top-Bio) reagent and flash froze in liquid nitrogen. We then proceeded to perform a standard phenol-chloroform extraction procedure for RNA isolation. RNA precipitation was performed overnight at 4°C by adding isopropanol to the lysis supernatant. The pellet was washed in 70% ethanol and dried at room temperature before being resuspended in H20.

Poly(A) RNAs were isolated from 20μg of total RNAs using a Dynabeads mRNA Purification Kit from Thermo Fisher Scientific by following the manufacturer’s protocol.

### Direct-cDNA library preparation

Sequencing libraries were prepared by following the SQK-DCS109 protocol provided on ONT’s website (https://nanoporetech.com) and by using the reagents provided in their direct-cDNA sequencing kit, along with the recommended 3rd party reagents.

Strand-switching reactions were performed using the Strand Switching Primer provided in the kit (SSP experiments), or by either replacing it with a SL1-specific probe (SL1 experiment) or omitting it from the reaction (No Primer experiments) (see Fig.1d).

### Data acquisiton and basecalling

All of the sequencing runs were performed using a MinION Mk1B sequencer connected to a computer running the MinKNOW software (version 2.0) with live-basecalling disabled. Following library preparation, the sample was directly loaded onto a MinION flow cell (R9.4 chemistry) accordingly to the manufacturer’s guidelines. Upon starting a run, we controlled the potency of each flow cell by making sure that a sufficient number of pores were actives and the experiments were run until we noticed a decrease of activity by monitoring channels state and the general output via the integrated dashboard. Flow cells were then washed and stored for further runs. In order to avoid cross-contaminations, flow cells were only re-used between duplicate experiments. In such case, we also adjusted the starting voltage because of voltage drift.

Basecalling of the obtained raw data (fast5 format) into readable files (fasta format) was performed using Guppy (version 2.3.7) with the flip-flop algorithm for better accuracy (options: c dna_r9.4.1_450bps_fast.cfg trim_strategy 0 qscore_filtering 1 calib_detect 1). Additionally, all of the basecalling step was performed on virtual machines running on Google cloud Platform and associated with an NVIDIA Tesla V100 GPU in order to enable guppy’s GPU-basecalling (options: x auto num_callers 16 chunks_per_runner 768 --chunk_size 500 gpu_runners_per_devices 8).

### Read alignment and processing

Following basecalling, reads were mapped with minimap2 v2.16 against *C. elegans* reference genome and transcriptome (WBcel235/ce11 genome assembly) obtained from the wormbase release WS270 (https://wormbase.org/).

Genomic alignments were generated by mapping the obtained cDNA sequences against *C. elegans* chromosomal sequences using the splice-aware alignment preset in order to account for the presence of intronic sequences in the genome (command: minimap2 ax splice). Furthermore, transcriptomic alignments were generated by using the messengers RNAs sequences as the reference along with minimap2 preset for mapping Oxford nanopore reads (command: minimap2 ax mapont).

The resulting alignment files, obtained in SAM format, were then sorted, indexed and converted to BAM format using SAMtools^30^.

### Genome browser visualisations

Genomic alignments were visualized using the Integrated Genome Viewer (IGV)^31^.

### Measure of strand bias

The measure of strand bias in each experiment was performed by using a custom python script which parsed all of the transcriptomic alignments and detected wether the read had been reversed during the alignment step or not. Since the reference sequences are in the 5’ to 3’ orientation, independently from the gene orientation, it is therefore possible to detect the strand origin of any given read by checking if the read had been reversed during mapping (antisense strand) or not (sense strand).

### Base quality assessment

When extracting base-quality from a single Nanopore read, we observed a lot of intrinsic variability. Therefore, in order to be able to compare the quality of the 5’ soft clipped region (unaligned) with that of the aligned portion of the reads, we used the following method: 1) we randomly selected a large number of reads 2) extracted their base-quality 3) determined the position of each base, relative to the alignment’s start 4) computed the mean base-quality for each position 5) plotted the average base-quality obtained. A visual representation of the method is shown in Sup. Fig. 6.

### Identification of Splice Leader sequences

Due to the nature of trans-splicing, we reasoned that a full-length cDNA read would map onto its transcript reference from the trans-splicing site to the polyadenylation site, leaving an unmapped (soft-clipped) region on each end, one in 5’ (the splice leader sequence) and one in 3’ (the poly(A) sequence). Hence, we decided to search for the splice leader sequence directly upstream of the alignment start. Moreover, in order to account for the inherent noise of Nanopore reads, and considering the small size of the SL motif (∼23nt), we decided to add some tolerance in order to allow for the detection of closely related sequences.

The method is briefly described as follows: 1) we parsed transcriptomic alignments and extracted the sequence (up to 100nt) directly upstream the alignment start 2) for each annotated SL sequence, we performed a semi-global alignment onto the extracted sequence 3) the alignment score was then evaluated using a custom threshold 4) if the score is equal or higher than the threshold, the SL is considered detected and we start evaluating another SL sequence, otherwise the sequence is shortened (2nt in 5’ are trimmed from the sequence) and we repeat the previous steps until we reach a truncated SL sequence of 7nt, in which case the SL is considered undetected 4) After evaluating every SL sequence, we then retrieved the SL sequence which scored the best overall. Reads for which we detected both a SL1 and a SL2 sequence are considered SL.X and reads for which we detected several SL2 sequences are considered SL2.X

### Measure of Splice Leader usage

To produce a statistically relevant analysis, we decided to measure SL ratio solely based on confident SL matches. Therefore, for any gene analyzed, we considered exclusively matches with an alignment score superior to 9 or in the direct vicinity of their alignment start (distance of 2nt or less). Doing so allowed us to include any match resulting from the detection of poorly sequenced SL motifs (due to a drop of base-quality as seen in Figure 1b) but for which our confidence in the result is strengthen by their presence near the alignment start, where splice leader sequences are expected to be found. This is supported by the fact that high-scoring matches (>15) are preferentially found just upstream the alignment start whereas low-scoring (<10) can be found all across the 5’ soft-clip sequence, but also with a noticeably higher frequency directly upstream their alignment start (Sup. Fig. 8).

Confident SL matches were then used to measure the ratio of SL2 to SL1 reads for a given gene or a given position.

### Identification of 5’ endogenous hairpins

To identify the presence of a stem-loop we examined the complementarity between two regions of the same read using the following method: 1) we selected reads originating from the antisense strand of the cDNA and for which no SL had been detected 2) we extracted the 5’ unaligned region corresponding to the potential complementary stem (positions -13 to +2 relative to alignment start). 3) the 5’ unaligned region extracted was then reverse complemented and mapped against the aligned region. 4) If at least 12nt (out of 15) were successfully mapped, we validated the presence of an endogenous hairpin in the read.

### Visualization of secondary RNA structures and hairpin motifs

We used the RNAfold web server (at http://rna.tbi.univie.ac.at/cgi-bin/RNAWebSuite/RNAfold.cgi) in order to generate secondary RNA structures based on SL or 5’ endogenous hairpins sequences. Additionally, we use the RNAbows web server (at http://rna.williams.edu/rnabows) to compute the probability of base-pairing between regions of the same read.

## Data availability

The direct-cDNA datasets used in this study have been deposited in the Sequence Read Archive (SRA) under the following accession code: PRJNA822363. and listed in the Supplementary Table 3.

The dataset table, generated by processing all of the obtained alignment files, and used in all computational analysis has been deposited on Figshare, along with all the other files generated as part of our analysis (DOI: 10.6084/m9.figshare.19131260).

A companion web-app, allowing the reader to view our results regarding any of the genes found in our sequencing experiments is directly available here: https://share.streamlit.io/florianbrnrd/elegans-trans-splicing/main/app/streamlit_app.py

## Code availability

The different scripts and methods used to perform the analysis were deposited on the following Github repository: https://github.com/florianbrnrd/elegans-trans-splicing.

## Acknowledgements

F.B was funded by IDEX Bordeaux International Doctorate Program, ISF grant 1339/17 and Sandwich scholarship from the Council of Higher Education. O.R is grateful to funding from the Eric and Wendy Schmidt Fund for Strategic Innovation (Polymath Award #0140001000) and funding from the Khan foundation.

## FIGURE LEGENDS

**Fig. 1: *C. elegans* messengers strand bias in Nanopore direct cDNA sequencing. a** Typical genome browser (IGV) view of direct cDNA reads aligned on *C. elegans* gene *lys-1*. Aligned bases from the sense strand reads are shown in pink and aligned bases from the antisense strands in purple unaligned soft clip region is shown with mismatches colored according to the observed base. Inset: overall strand bias observed across all detected transcripts. **b** Base quality measured in 5’ soft-clip and primary alignment. We measured the average base quality value over one million individual Nanopore reads. **c** Typical alignment of reads at the interface between the transcript match and the long soft-clipped region. The first bases of the soft-clip commonly correspond to the SL1 sequence followed by a partial antisense SL1 segment (blue to orange arcs) consistent with the extension of one of the endogenous hairpins represented as insets. The rainbow plot indicates base pairing between the 5’ soft clip and the trans spliced antisense strand. **d** Schematic representation of molecular events during the second strand synthesis step in various conditions of direct cDNA sequencing library preparation. Bottom diagram indicates the observed strand bias measured after Minion sequencing. **e** Schematic representation of the progression of a hairpin cDNA substrate during Nanopore sequencing. The helicase activity of the motor protein maintains a steady rate of transfer for the antisense strand while unwinding the double stranded cDNA. For the second strand the absence of double helix on the input side and/or the kinetics of the base pairing on the other side of the pore affects the signal quality and prevent accurate base calling of the sense strand.

**Fig. 2: Spliced leaders usage frequency. a** Genomic location of all *C. elegans sls* genes and structure of their 5’ hairpin strand. **b** Quantification of reads with high quality splice leader sequence reveals the usage frequency of each of the *sls* genes. SL2.x indicates reads for which the specific SL2 genes could not be unambiguously identified.

**Fig. 3: Quantitative analysis of SL sequence selectivity. a** bottom: A scatter-plot of the ratio of SL2/SL1 reads found against the distance to the closest upstream gene. Position with at least a 60% bias are colored red for SL2 and blue for SL1. Top: Distribution SL1 and SL2 biased positions confirms previous observations of a strong preference for SL2 splicing to occur at a distance of ∼100 nt downstream from the previous exon. **b** For SL2 trans spliced site represented in a (with more than 10 individual reads with an unambiguous SL2 sequence) we represent the frequency of usage of each SL2. Top: Genes located around 100 nt downstream of another coding gene. Bottom: Genes located further away from the closest upstream coding gene with genes predominantly SL2 on the left and predominantly SL1 on the right.

**Fig. 4: Nature of non-trans-spliced messengers a** Gene expression and trans splicing status. We grouped gene in the *C. elegans* genome for which a direct cDNA read was obtained in this study according to the presence of evidence of trans splicing in this study and/or in the compendium of RNA-seq we previously analysed (Tourasse, 2017). For each gene we plotted the number of Nanopore reads we obtained as a proxy for gene expression level. **b** Trans splicing detection level. For each gene we identified the most frequently detected 5’ alignment starting position and plotted the number of reads at this position vs the number of those reads that for which a Spliced Leader sequence could be detected. Genes for which the most represented start position had at least 100 reads but had less than 10% SL detection are represented in green. **c** Poorly trans spliced mRNA have a propensity to form a 5’ stem loop structure. For the three genes highlighted in panel b we show a partial alignment of the end of the aligned region and the beginning of the soft clipped region.

**Fig. 5: Quantitative representation of trans splicing prevalence. a** We randomly selected 250,000 reads from each category and plotted the average Q-score around the end of the alignment start position. **b** Repartition of reads for which a trans splicing event could be confirmed (SL), with terminal self complementarity (hairpin) or with unknown 5’ status. **c** Schematic representation of the trans-splicing information recovered in this work. Each alignment start position observed was plotted at the corresponding genomic position with the number of supporting reads. The dots are colored according to the legend on the right. With red indicating a majority of SL reads, green a majority of endogenous hairpin reads and blue reads with no evidence for either. Dotted vertical lines indicate the position of a putative start codon. *nlp-36* displayed a single SL1 trans-spliced site. M60.4 presents two trans-splicing sites indicative of two alternative promoters with distinct SL1/2 ratios. The Y37E3.7/Y37E3.8 operon with typical SL1 and SL2 preference. *lev-11* gene distal promoter carries an endogenous hairpin while the proximal promoter is mainly SL1.

**Fig. 6: Three types of *C. elegans* mRNA 5’ extremities**. mRNAs derived from gene adjacent to their promoters receive mostly SL1. SL2 is preferentially trans-spliced to genes located ∼100nt downstream of another gene. Non-trans-spliced mRNAs have 5’ self-complementarity that mimics the structure of SLs.

## SUPPLEMENTARY FIGURES

**Sup. Fig. 1:**
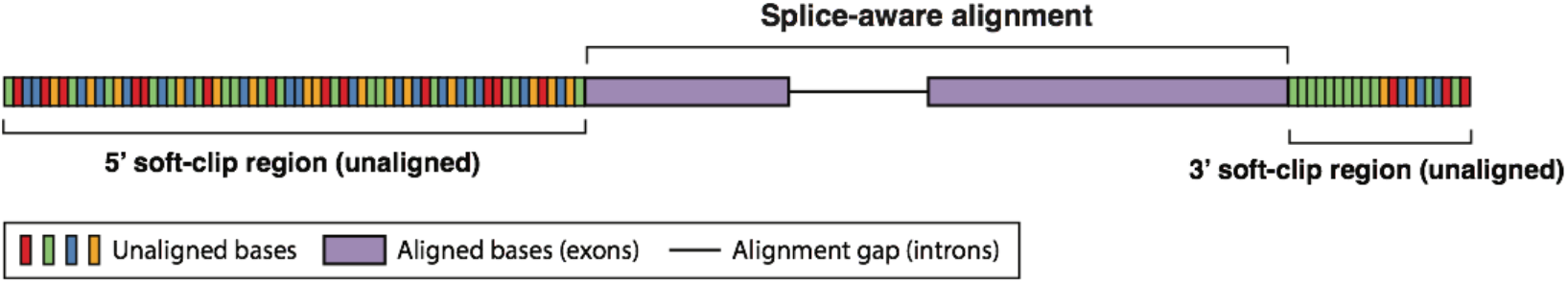
Schematic representation of genomic read alignments obtained in *C. elegans* direct-cDNA experiments. Purple region represent the region of a read mapped onto the genome. Soft-clip regions (in 5’ and 3’, per gene orientation) corresponds to unmapped part of the read, typically containing motifs from sequencing adapters and exogenous RNA sequences such as Spliced Leader (SL) and polyadenylation motif. In our experiments, 5’ soft-clips region are unexpectedly long.

**Sup. Fig. 2:**
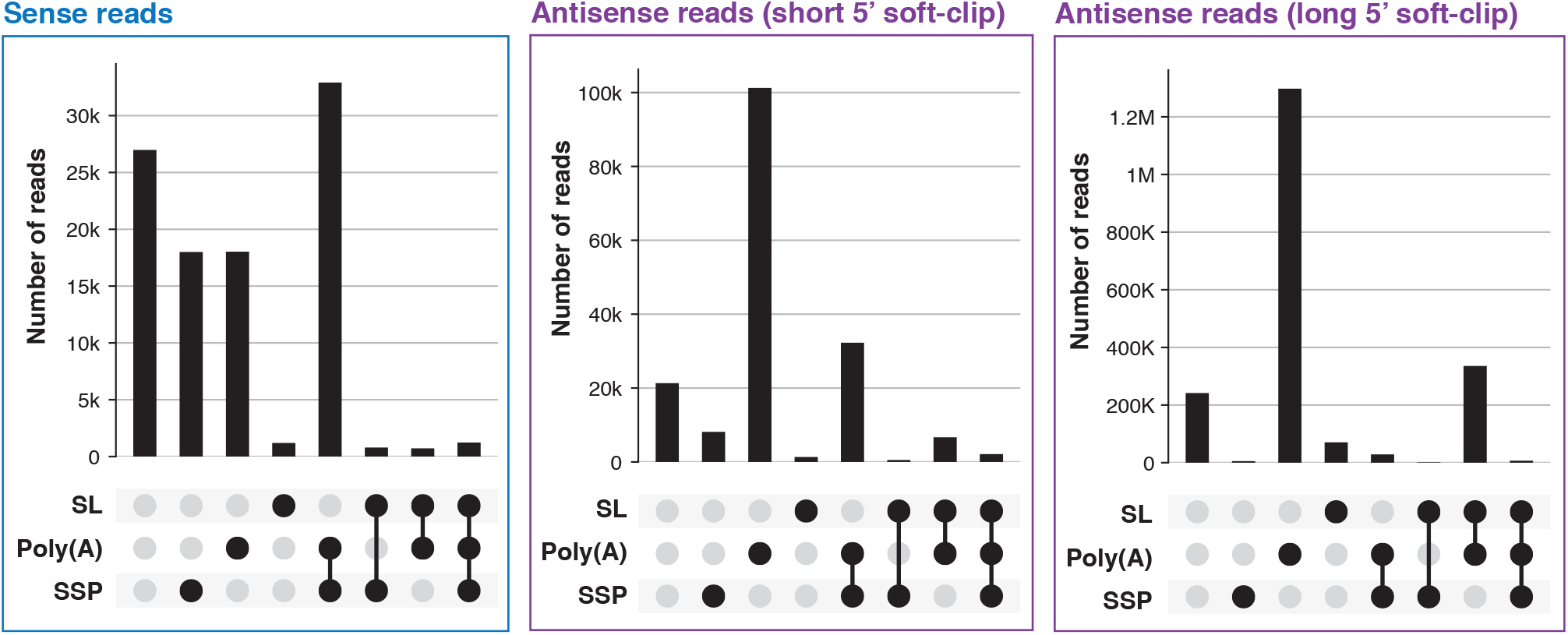
Identification of direct-cDNA sequencing adapters in reads with short soft-clips regions. For each read evaluated, we performed a sequence search for the SSP and SL motifs on the 5’ soft-clip region, and for the poly(A) motif on the 3’ soft-clip region.

**Sup. Fig. 3:**
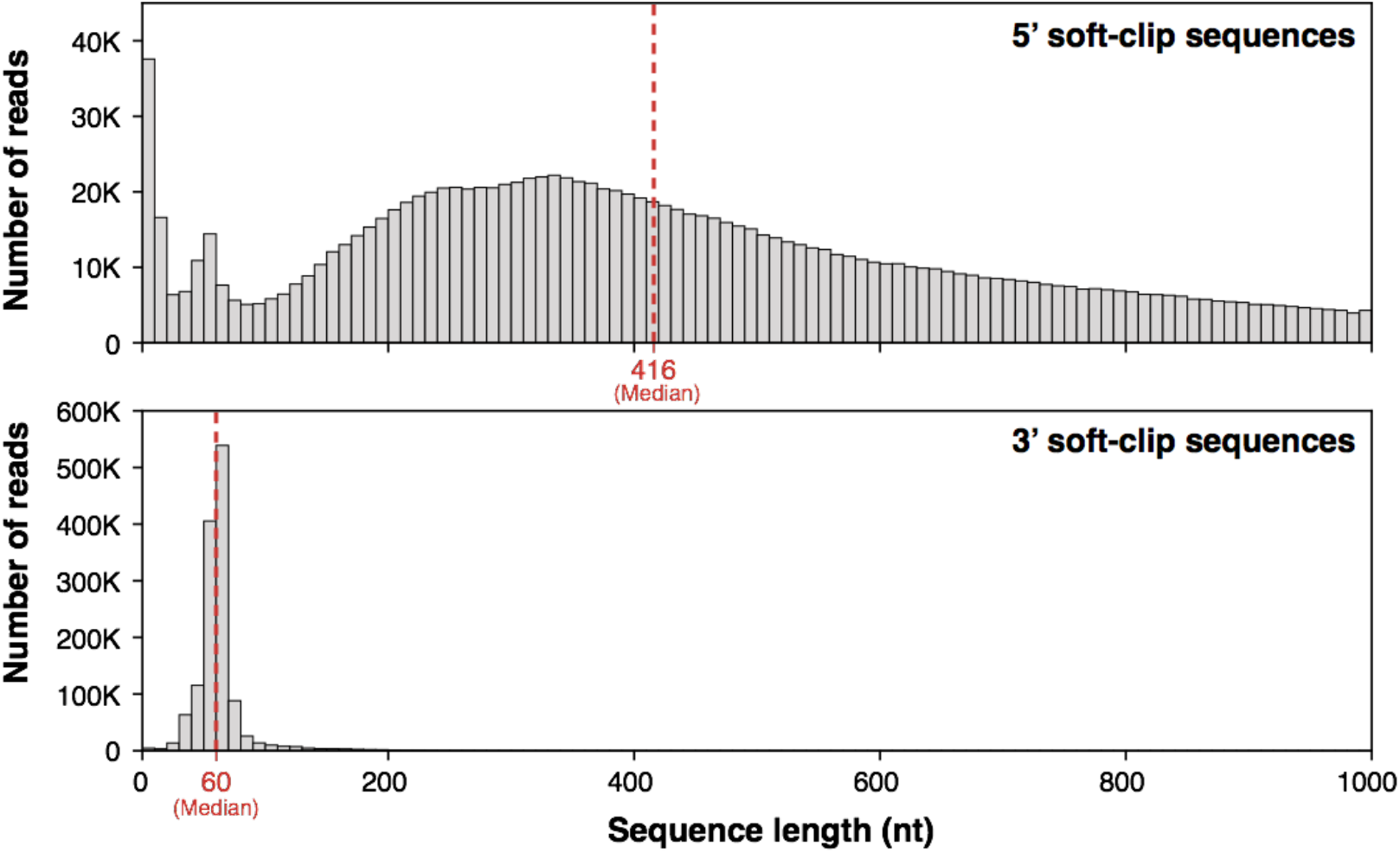
Measured length of 5’ and 3’ soft-clip. Length of each region was plotted as a histogram (bins of 10nt) and for each, median value was measured (shown in red). **Top:** 5’ soft-clip sequences (median length = 416nt). **Bottom:** 3’ soft-clip sequences (median length = 60nt).

**Sup. Fig. 4a:**
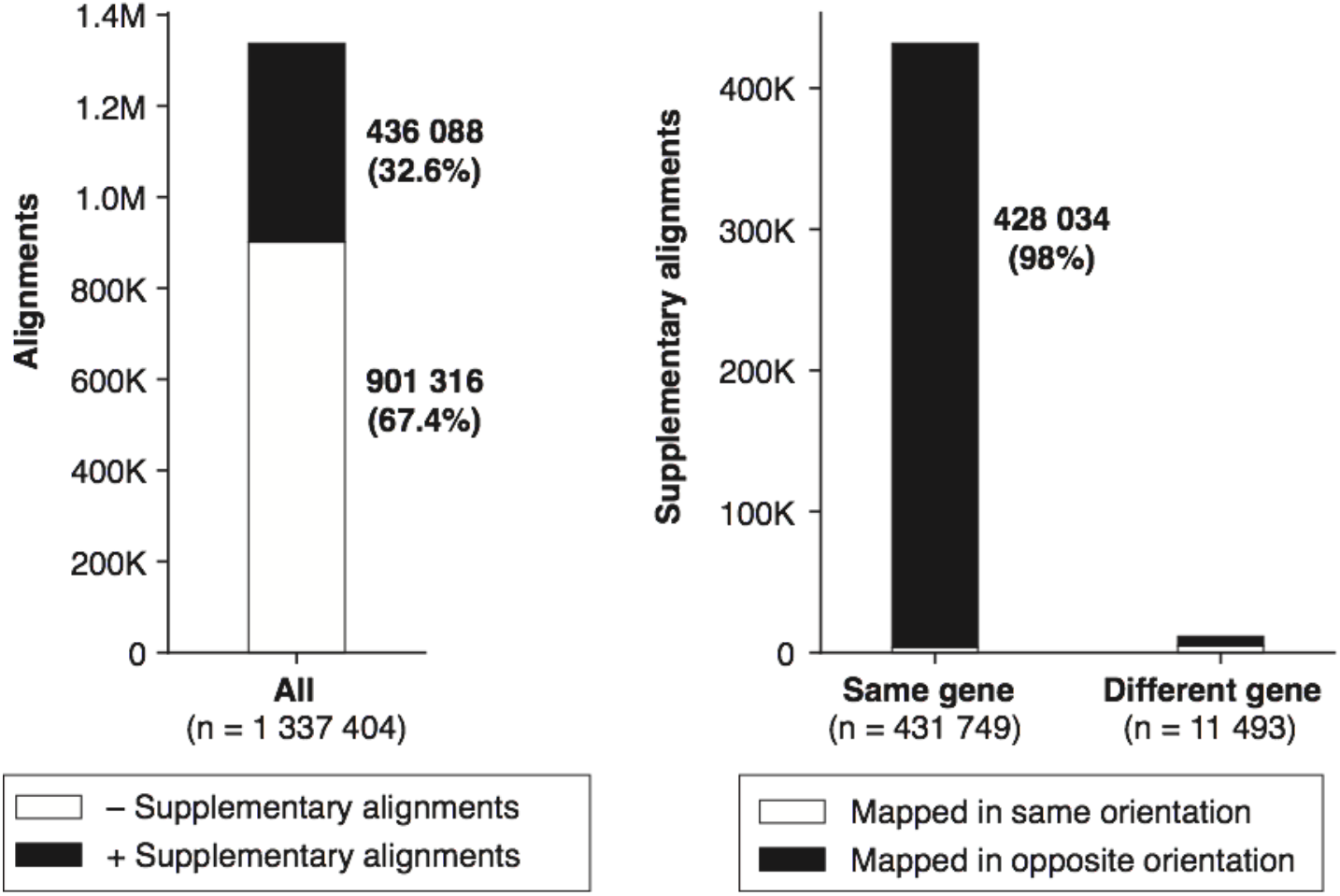
Origin of supplementary alignments. 32.6% of reads with primary alignments possess supplementary alignments. Among those, 98% are mapping to the same gene than the primary alignment, but in the opposite orientation.

**Sup. Fig. 4b:**
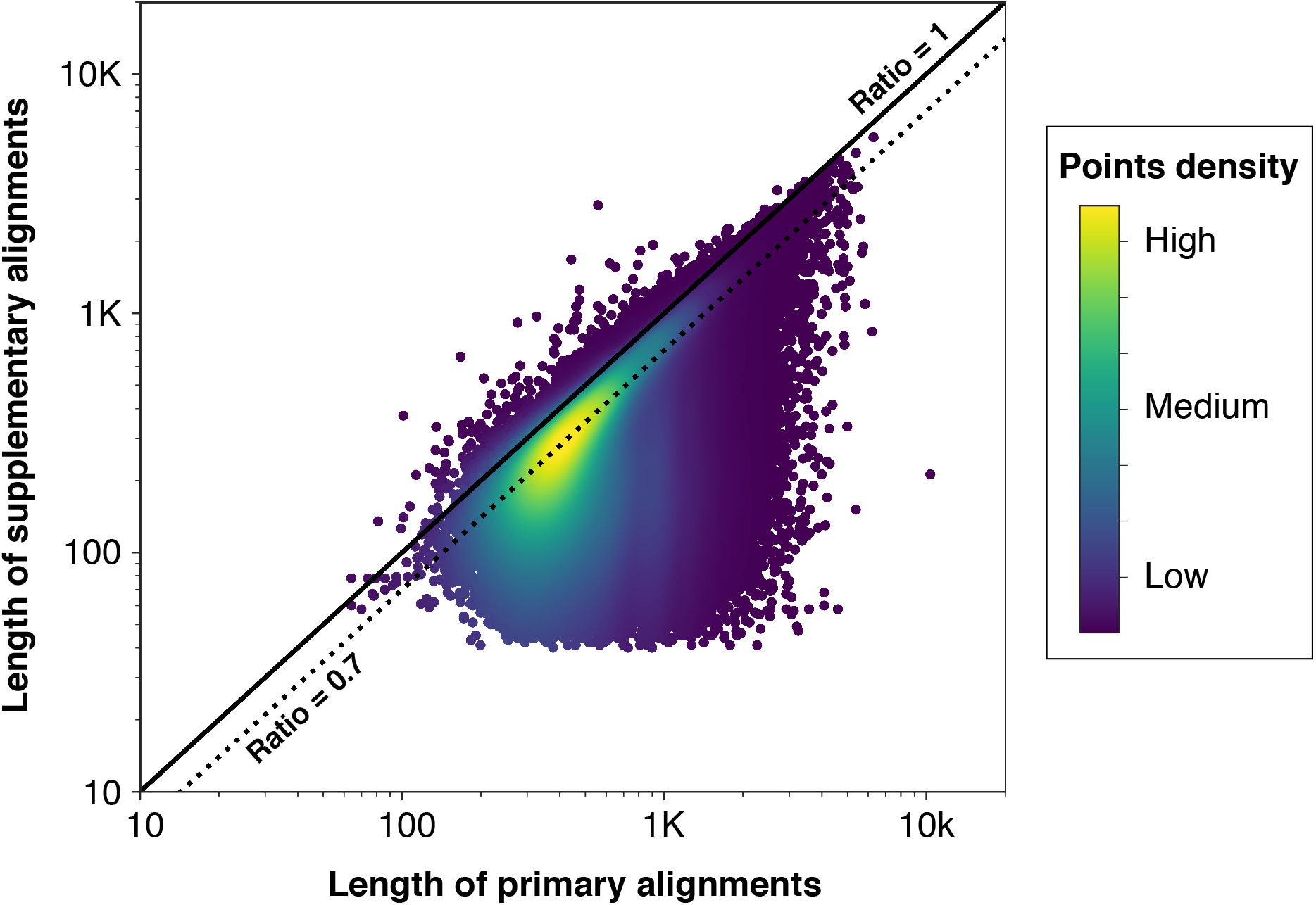
Size distribution of supplementary alignments. Sizes of primary and supplementary alignments were compared for reads possessing both. Solid black line represent supplementary alignments that are the same size as their primary alignment (ratio 1/1). Dotted black line represent supplementary alignments that 70% of the size of their primary alignment (ratio 1/0.7).

**Sup. Fig. 5:**
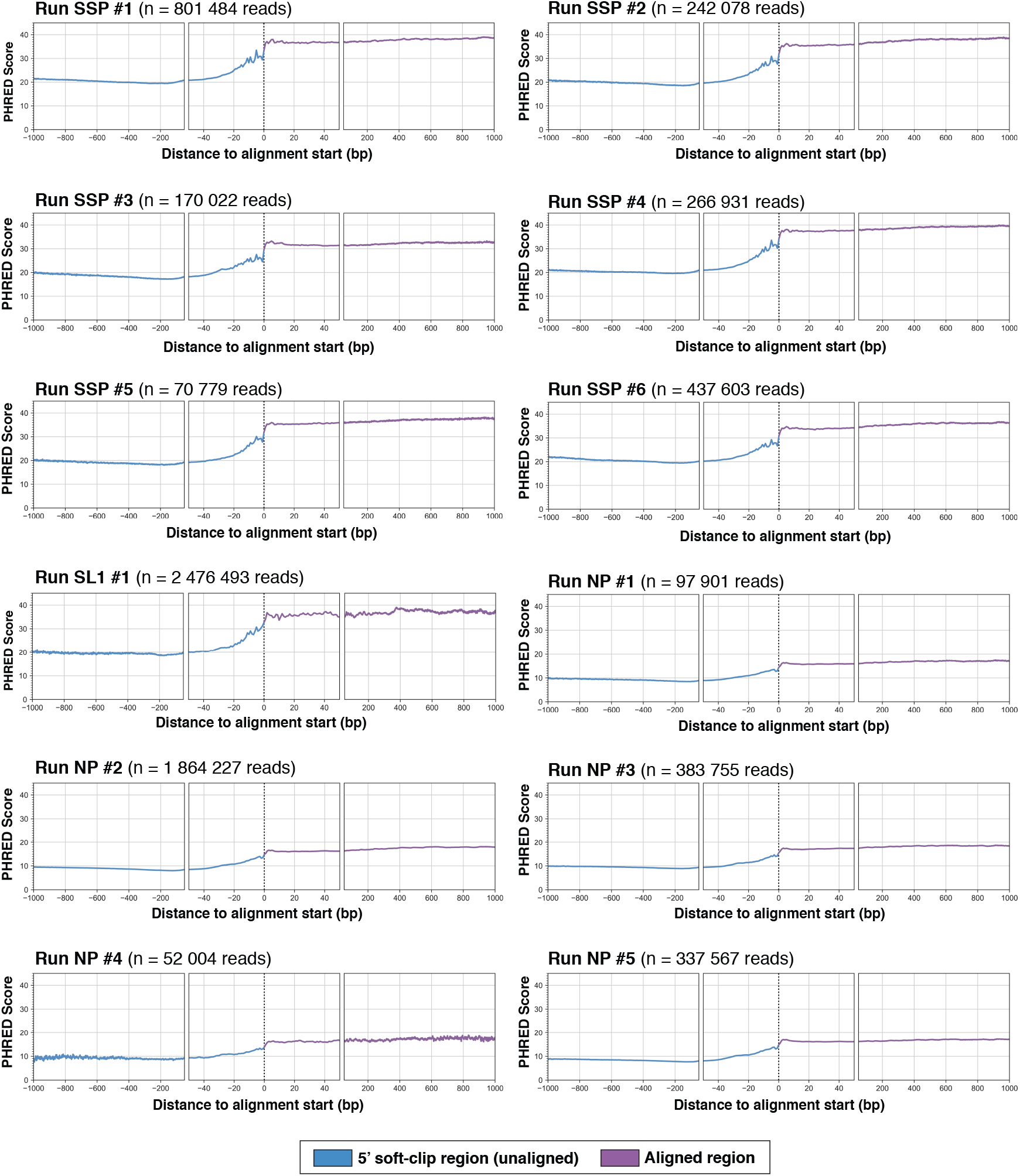
Base quality measures in each experiment. We measured the average base quality value over all the Nanopore reads obtained in each sequencing experiment. 5’ soft-clip region is represented in blue and aligned region (primary alignment) is represented in purple.

**Sup. Fig. 6:**
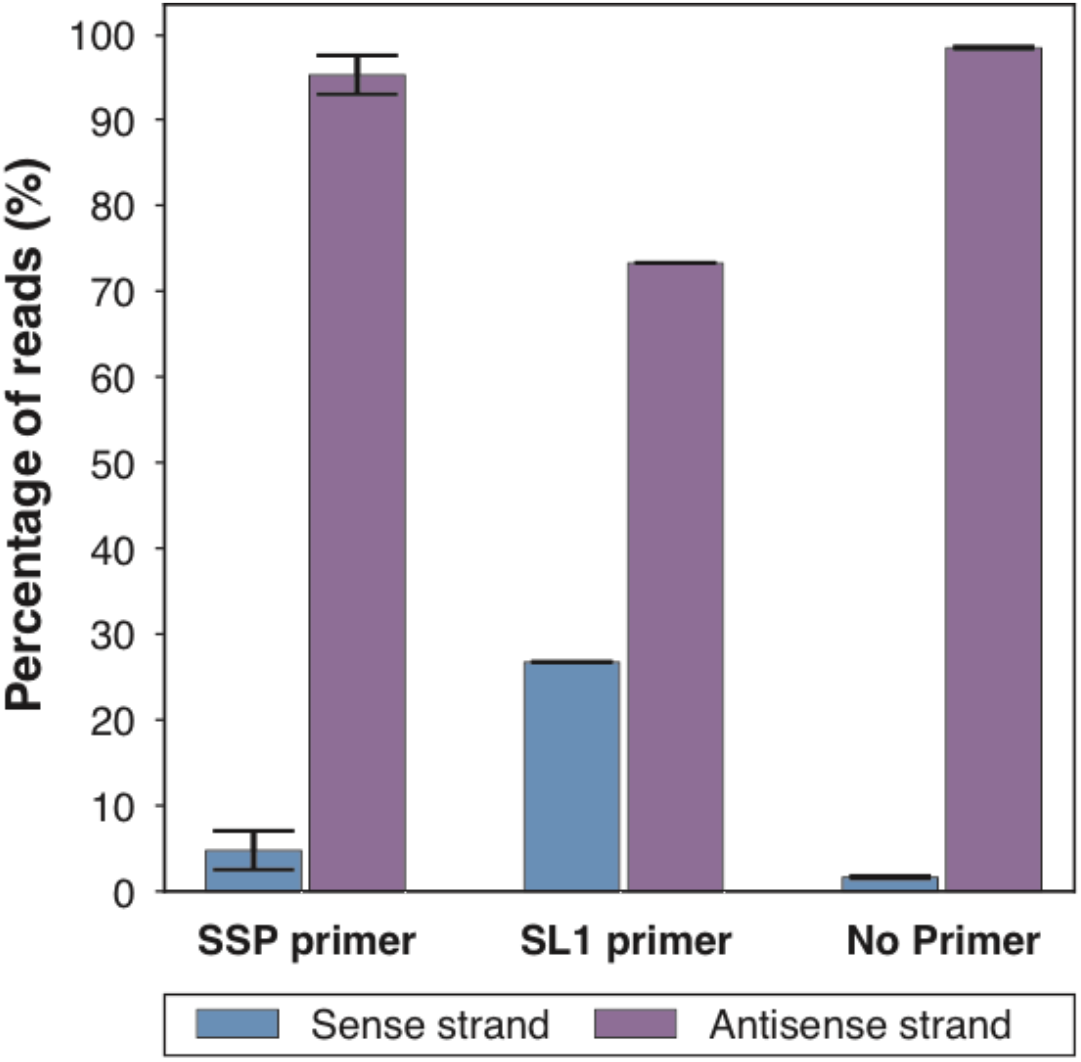
Strand bias in ONT direct-cDNA experiments. For each experiment, we measured the number of reads coming from each strand of the cDNA molecule sequenced. **SSP primer:** 2nd strand synthesis using the SSP primer (n=6). **SL1 primer:** 2nd strand synthesis using a SL1-specific primer (n=1). **No primer:** 2nd strand synthesis without using any primer (n=5).

**Sup. Fig. 7:**
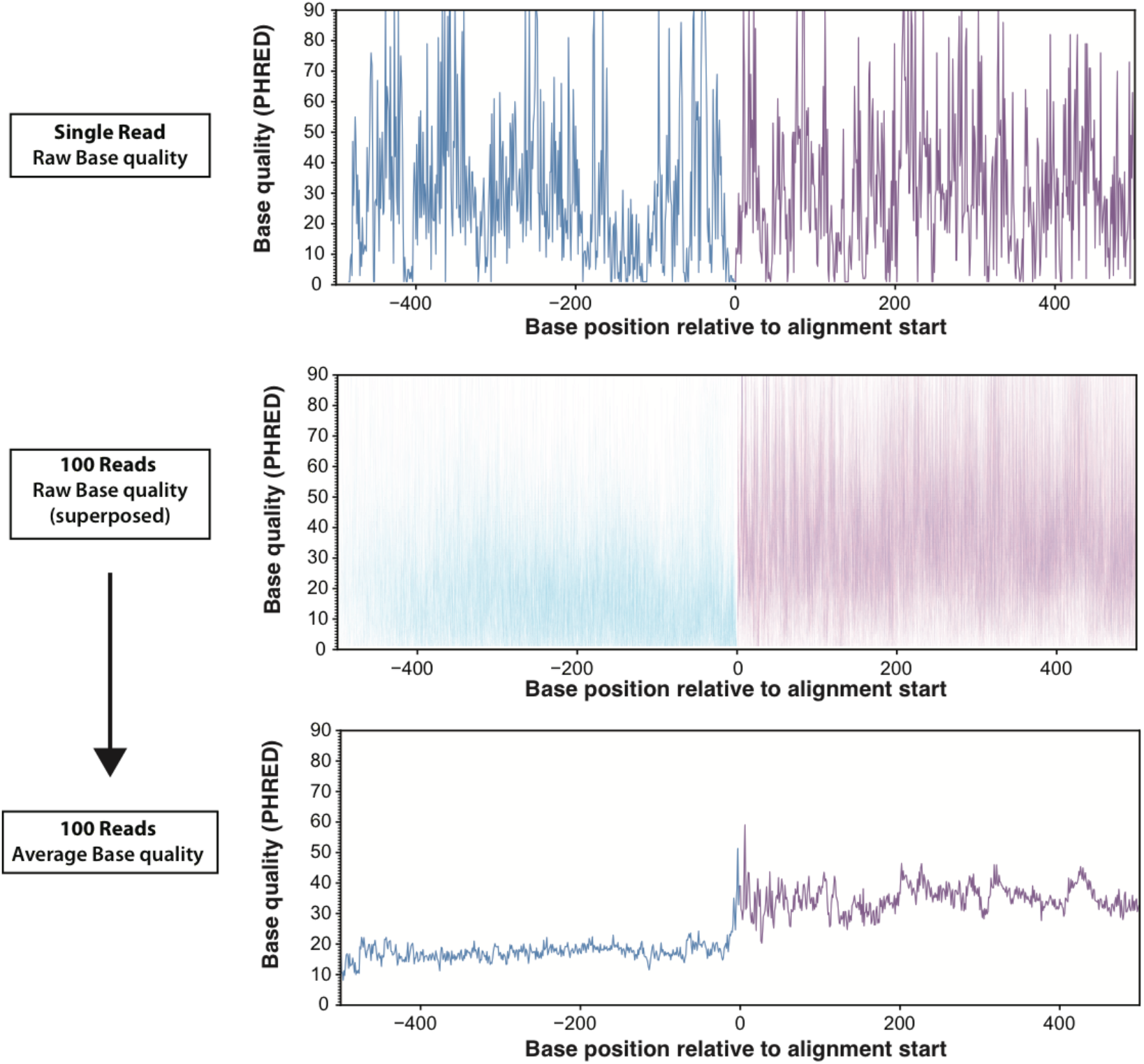
Method for evaluating base quality. Single read’s base quality contains too much variability for accurate measurement (top panel). Trends in base quality can be observed when looking at larger samples of reads (middle panel - 100 reads shown together). Averaging the base quality per position across a large sample of reads (bottom panel) allows to accurately visualize differences in base quality between different regions of the reads. Unaligned 5’ soft-clip region is shown in blue and alignment region in purple.

**Sup. Fig. 8:**
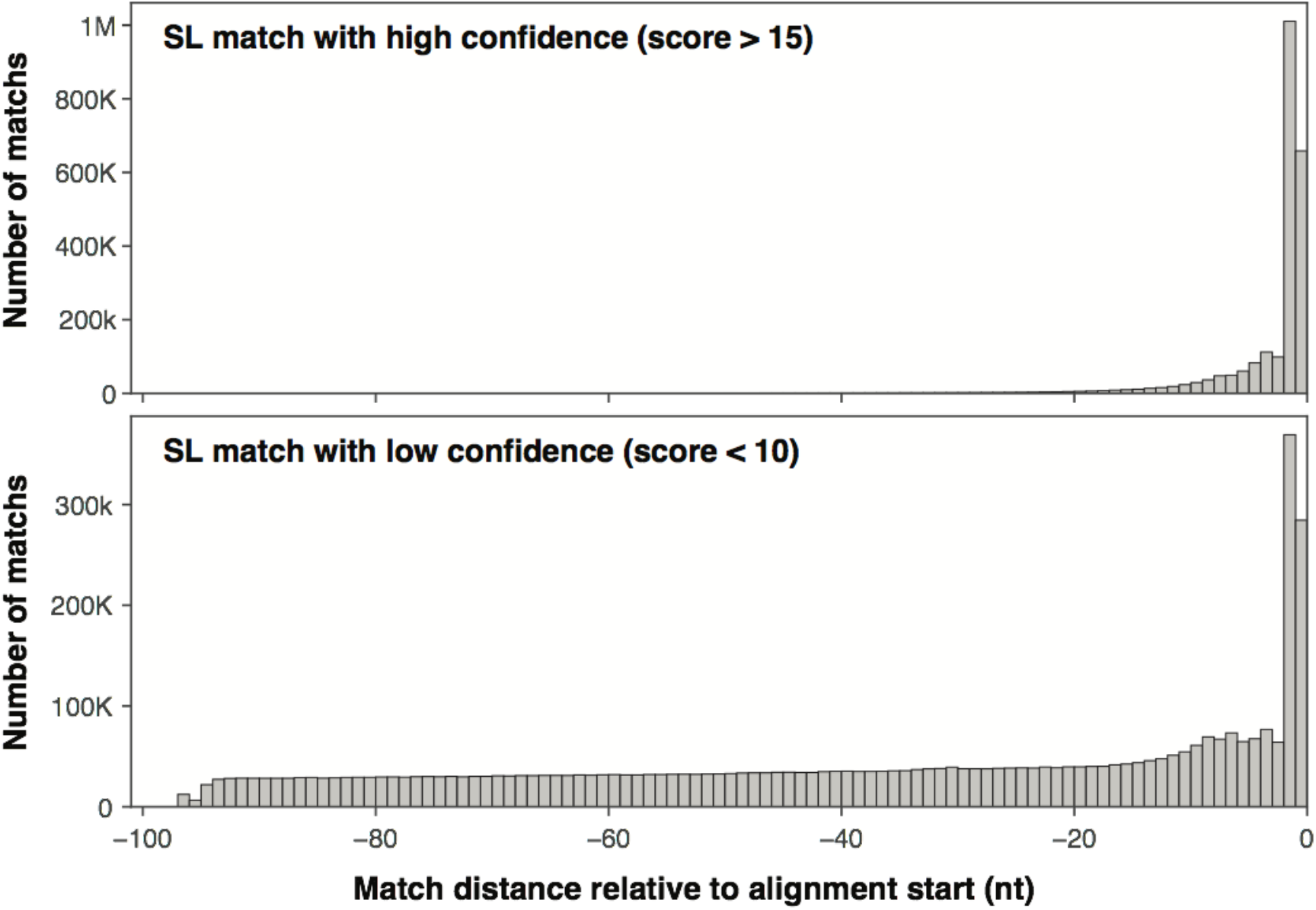
High confidence SL matches are preferentially located near the alignment start. For each SL sequence identified, we measured their distance to the alignment start. As expected, matches which scored high (top panel) are found in the direct vicinity of the alignment start. Low-scoring matches (bottom panel) are generally found all across the 5’ soft-clip region, but a sub-set of those are found where expected making those more reliable.

